# Nitrogen cost minimization is promoted by structural changes in the transcriptome of N deprived *Prochlorococcus* cells

**DOI:** 10.1101/087643

**Authors:** Robert W. Read, Paul M. Berube, Steven J. Biller, Iva Neveux, Andres Cubillos-Ruiz, Sallie W. Chisholm, Joseph J. Grzymski

**Affiliations:** Division of Earth and Ecosystem Sciences, Desert Research Institute, Reno, NV, USA; Department of Civil and Environmental Engineering, Massachusetts Institute of Technology, Cambridge, MA, USA; Microbiology Graduate Program, Massachusetts Institute of Technology, Cambridge, MA, USA; Department of Biology, Massachusetts Institute of Technology, Cambridge, MA, USA

## Abstract

*Prochlorococcus* is a globally abundant marine cyanobacterium with many adaptations that reduce cellular nutrient requirements, facilitating growth in its nutrient-poor environment. One such genomic adaptation is the preferential utilization of amino acids containing fewer N-atoms, which minimizes cellular nitrogen requirements. We predicted that transcriptional regulation might be used to further reduce cellular N budgets during transient N limitation. To explore this, we compared transcription start sites (TSSs) in *Prochlorococcus* MED4 under N-deprived and N-replete conditions. Of 64 genes with primary and internal TSSs in both conditions, N-deprived cells initiated transcription downstream of primary TSSs more frequently than N-replete cells. Additionally, 117 genes with only an internal TSS demonstrated increased internal transcription under N-deprivation. These shortened transcripts encode predicted proteins with ~5-20% less N content compared to full-length transcripts. We hypothesized that low translation rates, which afford greater control over protein abundances, would be beneficial to relatively slow-growing organisms like *Prochlorococcus*. Consistent with this idea, we found that *Prochlorococcus* exhibits greater usage of glycine-glycine motifs, which cause translational pausing, when compared to faster growing microbes. Our findings indicate that structural changes occur within the *Prochlorococcus* MED4 transcriptome during N-deprivation, potentially altering the size and structure of proteins expressed under nutrient limitation.

## Introduction

Primary productivity is limited by nitrogen (N) availability in many ocean ecosystems (Tyrrell, 1999; Deutsch *et al.*, 2007; Moore *et al.*, 2013), and organisms that live there have adaptations that help them survive in low nutrient conditions. These adaptations include small cell sizes, which facilitate nutrient transport by increasing surface:volume ratios and reducing the absolute cellular requirement for nutrients (Munk and Riley, 1952; Gavis, 1976; Chisholm, 1992), as well as small genomes and a proteome that is N cost minimized (Grzymski and Dussaq, 2012). The concept of cost minimization was originally based on an observation that assimilatory proteins for a specific element contain relatively less of that specific element than average proteins in the same organism (Baudouin-Cornu *et al.*, 2001). For many oligotrophic microorganisms, however, cost minimization extends beyond the proteins involved in assimilation and is observed across the proteome; e.g. the genomes of N cost minimized oligotrophic microbes code for proteins that are, on average, reduced in amino acids containing added N side chains as compared to their coastal counterparts (Grzymski and Dussaq, 2012). As an evolutionary trade-off, the proteomes of N-cost minimized organisms typically have slightly larger mass or more C atoms – an element which is not usually limiting for autotrophs (Grzymski and Dussaq, 2012).

The marine cyanobacterium *Prochlorococcus* MED4 is a striking example of a N-cost minimized organism (Grzymski and Dussaq, 2012). *Prochlorococcus* is often the numerically dominant primary producer in oligotrophic waters (Flombaum *et al.*, 2013), and can be broadly divided into two main subgroups, made up of high-light adapted and low-light adapted cells. High-light adapted *Prochlorococcus* cells are the smallest cyanobacterial cells in the oceans, and also have the smallest genomes of any free-living photosynthetic cell. The abundance of high-light adapted *Prochlorococcus* cells typically exceeds that of low-light adapted cells by several orders of magnitude in surface waters (Ahlgren *et al.*, 2006; Zinser *et al.*, 2006; 2007; Malmstrom *et al.*, 2010). The divergence of high-light adapted *Prochlorococcus* from their low-light adapted relatives is correlated with a reduction in the overall G+C content of their genomes, thereby reducing the N requirements of these cells due to the bias of low G+C codons to encode low N-containing amino acids (Grzymski and Dussaq, 2012).

In addition to reducing the N content of their proteomes, high-light adapted *Prochlorococcus* have undergone a process of genome streamlining in which the number of coding sequences has been reduced, in part, by dispensing with most regulatory proteins (Grzymski and Dussaq, 2012; Giovannoni *et al.*, 2014). How, then, do these cells respond to changing environmental conditions? Recent studies have revealed that, in addition to traditional protein regulators (transcription factors, sigma factors, etc.), many bacteria also utilize RNA-based mechanisms to regulate transcript abundance (Sharma *et al.*, 2010). *Prochlorococcus*, like other streamlined microbes such as *Helicobacter pylori* , has an unexpectedly complex transcriptome for a small genome, that includes cis and trans regulating RNA, other non-coding RNAs, varying cistronic and polycistronic operon regulation, and mRNAs with short half-lives (Steglich *et al.*, 2010; Voigt *et al.*, 2014). These regulatory characteristics likely enable responses to environmental signals without a complex protein-based regulatory network.

Given the degree to which N cost minimization has been selected for in the genome of *Prochlorococcus* MED4 over evolutionary time, we wondered whether the transcriptional response in these cells has also been optimized to further reduce N requirements. We hypothesized that transcriptome changes could help provide additional N savings during nutrient limited growth – a concept we define as transcriptomic cost minimization. To test this hypothesis, we compared the primary transcriptomes (the set of unprocessed transcripts that allow experimental determination of transcription initiation sites) of *Prochlorococcus* MED4 grown under N-deprived and N-replete conditions, allowing us to examine alterations in the transcriptional start sites under N-deprivation. Our results uncovered responses to N stress that highlight the capacity of already ‘streamlined’ cells to dynamically minimize their N requirements in response to N-deprivation.

## Materials and Methods

### Cell cultures

Axenic *Prochlorococcus* MED4 cells were grown in batch culture at 25 °C under continuous illumination of ~55 µmol photons m^−2^ s^−1^ using cool, white, fluorescent bulbs. Cultures were grown in Pro99 media (Moore *et al.*, 2007), prepared with 0.2 µm filtered surface seawater from the South Pacific Subtropical Gyre (26.25 °S, 104 °W) and amended with 3.75 mM TAPS buffer (pH=8) and 6 mM sodium bicarbonate to control pH. Cells were allowed to acclimate to these temperature, light and media conditions by successively transferring the cultures four times during mid-exponential phase (~20 generations).

Triplicate 4.5 L batch cultures were grown in 10 L clear polycarbonate carboys. Bulk culture fluorescence was measured daily using a 10AU fluorometer (Turner Designs, Sunnyvale, CA, USA). Cells were preserved for flow cytometry by fixation in 0.125% glutaraldehyde and storage at -80 °C. Once the cultures reached mid-exponential phase, 3.8 L of each culture was concentrated by centrifugation in four 1 L centrifuge bottles (Beckman Coulter, Brea, CA, USA) using a JLA-8.1000 rotor in an Avanti J-20 XP centrifuge (Beckman Coulter, Brea, CA, USA) at 25 °C and 9000 g (6857 rpm) for 15 min. Each cell pellet was washed once by re-suspension in 375 mL of N-depleted media (lacking ammonium chloride addition) and concentrated again by centrifugation. Half of the washed cells were re-suspended in 1.8 L of N-replete Pro99 media, and the remaining half of the washed cells were re-suspended in 1.8 L of N-depleted Pro99 media (lacking ammonium chloride addition).

Samples for bulk culture fluorescence, cell enumeration by flow cytometry, fluorescence induction measurements, and RNA extraction were collected at 0, 3, 12 and 24 hours following the re-suspension of cells in either nitrogen-replete or nitrogen-depleted media. At each time point, bulk culture fluorescence was measured using a 10AU fluorometer (Turner Designs, Sunnyvale, CA, USA), and cells were preserved for flow cytometry by fixation in 0.125% glutaraldehyde and storage at -80 °C. Following the experiment, fixed cells were enumerated using an Influx Cell Sorter (BD Biosciences, San Jose CA, USA), as previously described (Olson *et al.*, 1985; Cavender-Bares *et al.*, 1999). Photochemical conversion efficiency (Fv/Fm) was measured using a FIRe Fluorescence Induction and Relaxation System (Satlantic, Halifax, Nova Scotia, Canada); cells were acclimated in the dark for 30 min to relax photosynthetic reaction centers before fluorescence induction curves were obtained using single turnover flash (STF) with blue light (450 nm with 30 nm bandwith). Raw data were processed using fireworx (Dr. Audrey B. Barnett, Dalhousie University, Halifax, Nova Scotia, Canada) in MATLAB. Specific growth rates of replicates in the 4.5 L batch starter cultures were estimated from the log-linear portion of growth curves constructed from bulk culture fluorescence and cell counts obtained from flow cytometry analysis. Fluorescence-based measurements yielded a growth rate of 0.60 d^−1^ ± 0.01 (1 standard deviation) and flow cytometry based measurements yielded a growth rate of 0.67 d^−1^ ± 0.14 (1 standard deviation).

At each time point, biomass for RNA extraction was obtained by centrifugation of 250 mL of culture in 250 mL centrifuge bottles (Beckman Coulter, Brea, CA, USA) using a JA-14 rotor in an Avanti J-20 XP centrifuge (Beckman Coulter, Brea, CA, USA) at 25 °C and 9000 g (9666 rpm) for 15 min. Cell pellets were re-suspended in 750 µl of RNA Denaturation Solution (Ambion ToTALLY RNA Total RNA Isolation Kit, Life Technologies, Grand Island, NY, USA) by pipetting, frozen in liquid nitrogen and stored at -80 °C until processing, at which point total RNA was extracted using the Ambion ToTALLY RNA kit (Life Technologies, Grand Island, NY, USA), yielding ~12 µg of total RNA from each pellet.

### RNA sequencing overview

Total RNA from each sample was divided and used in two sequencing workflows: gene expression analysis using strand specific sequencing of terminator exonuclease (TEX)-treated RNA (all time points) and primary transcriptome analysis using 5’ rapid amplification of cDNA ends (RACE) tagRNA-seq (12 and 24 hour time points). For gene expression analysis, it is necessary to account for transcripts within the total RNA pool that have undergone triphosphate to monophosphate conversion for ultimate degradation (Celesnik *et al.*, 2007; Schoenberg, 2007). A caveat of doing time-series analysis of gene expression using total RNA is that these signals are not removed and changes can be obfuscated, especially if half-lives of RNA are pathway or gene-dependent – a likely scenario with *Prochlorococcus*. The half-life differences of RNAs (Steglich *et al.*, 2010) could play a significant role in interpreting differences in RNA treatment methodology. For these reasons, we chose to use TEX-treated RNA for gene expression analysis. For primary transcriptome analysis and determination of transcription start sites, 5’ RACE tagRNA-seq was used because it selectively captures 5’ triphosphate ends at single nucleotide resolution. This produces the sawtooth-like pattern observed at transcriptional start sites throughout the genome (Sharma *et al.*, 2010).

### Ion Torrent sequencing for gene expression analysis

RNA was treated with Terminator Exonuclease (Epicentre, Madison, WI, USA) to remove rRNA contamination and sequenced at the Nevada Genomics Center (Reno, NV, USA). Library preparation was performed using the Ion Total RNA-Seq Kit v2 (Life Technologies, Grand Island, NY, USA), which fragmented the total RNA and used reverse transcriptase to produce cDNA for strand-specific sequencing on the Ion platform. Strand specific Ion Torrent Sequencing (Life Technologies, Grand Island, NY, USA) yielded 130 bp reads. Raw sequencing reads were uploaded to the sequencing read archive (NCBI accession: SRP078366, SRX1939126), and were inspected using FastQC (Andrews, 2009) to determine quality, ambiguous read percentage and relative amount of sequence reads. Aliases for the individual run names can be found at https://trace.ddbj.nig.ac.jp/DRASearch/experiment?acc=SRX1066875. Ion-Torrent RNA sequencing resulted in an average of 2.62x10^7^ ± 5.77x10^6^ raw reads per cDNA library for the terminator exonuclease (TEX)-treated, N-deprived cells and 2.29x10^7^ ± 6.99x10^6^ reads for the TEX-treated, N-replete cells. The average quality score was Q24 (Supplementary Table S1).

Sequencing reads were utilized as the input for Rockhopper (McClure *et al.*, 2013) for bacterial RNA-sequencing analysis. Seed length was 15% of the read length, and any read with mismatches greater than 15% of the read was disregarded. This removed approximately 10% of the reads from each treatment. Differential expression was calculated against a negative binomial distribution between the experimental (N-deprived) cells and the control (N-replete cells) at each time point (0, 3, 12, 24 hours post N removal) using the DESeq algorithm (Anders and Huber, 2010; Tjaden, 2015). Significantly differentially abundant transcripts were determined as those with a *p*-value < 0.05 when comparing the two treatments. Pearson correlation coefficients to previous research (Tolonen *et al.*, 2006) were larger for genes in the top 50% of mean expression level; these differences are likely due to low read depth mapping in non-regulated genes. Therefore, expression changes are only reported for those transcripts with normalized Rockhopper expression values in the top 50% of the data set.

### Illumina sequencing for primary transcriptome analysis

To determine the primary transcriptome of MED4, we analyzed transcriptional mappings at 12 and 24 hours post N-deprivation relative to the N-replete controls. Total RNA was treated with Ribo-Zero rRNA Removal Kit for bacteria (Epicentre) to remove rRNA and any processed transcripts with 5’ monophosphate ends. The samples were next treated with 5' RNA polyphosphatase (5RPP; Epicentre) in order to convert the remaining 5' triphosphate structures into 5' monophosphate ends in preparation for adapter ligation. 5’ Illumina TruSeq adapters were ligated to the monophosphate groups of the transcripts and cDNA was synthesized. The tagged 5' cDNA fragments were then specifically amplified with PCR and sequenced using 5’ RACE tagRNA-seq on a NextSeq 500 system with 75 bp read lengths (vertis Biotechnologie AG). To visualize the sequencing reads (NCBI accession: SRP078366, SRX1939254) and transcriptional changes, reads were mapped to the *Prochlorococcus* MED4 genome using the segemehl short read aligner (Hoffmann *et al.*, 2009; 2014). Resulting sam files containing mapped reads were converted into sorted and indexed bam files using samtools (Li *et al.*, 2009). A single alignment run of Transcription Start Site Annotation Regime (TSSAR) software was performed to determine start site differences between the transcriptional start site enriched samples and non-enriched samples in the N-replete and N-depleted samples (Amman *et al.*, 2014). In keeping with the definitions considered by the TSSAR algorithm, we classified TSSs located on the opposite strand of an annotated gene as antisense; TSSs located within 250 nt upstream of the gene’s annotated TSS as primary start sites; and TSSs within the annotated gene as internal (Amman *et al.*, 2014). Orphan TSSs were those that did not fall into any of these three categories, and we noted that some TSSs can represent a combination of multiple categories. Transcription from internal start sites in the sense orientation yield mRNAs that are called intraRNAs. Parameters for determining start site changes were a p-value of 1x10^−10^, a noise threshold of 4 and a merge range of 5. All TSSs identified were hand-curated by uploading the sorted bam files into the Integrative Genomics Viewer, which placed the reads in the correct position and orientation in relation to the annotated *Prochlorococcus* MED4 genome (Robinson *et al.*, 2011; Thorvaldsdóttir *et al.*, 2013). Positions with a read count difference of less than 100 between the enriched and non-enriched samples were ignored. The TSSs identified by TSSAR after 24-hour post N-depletion were used to determine which TSSs were conserved between 12 and 24 hours post N-depletion. Sorted bam files were opened in R and read coverage figures were produced using the ggplot2 package (Wickham, 2009). If both a primary and internal TSSs were present, the abundance of transcripts derived from an internal TSS was calculated by comparing the read mapping at the internal TSS to the read mapping at the primary TSS for both the N-depleted and N-replete samples. The magnitude of internal transcription in each treatment was then compared against each other. Only internal TSSs resulting in transcript abundance greater than 10% compared to their primary TSS were used. If no primary TSS was present, transcription from internal TSS was calculated by directly comparing read mapping in the N-depleted sample to read mapping in the N-replete sample.

### Multiple sequence alignment and protein threading

For several genes with possible NtcA binding sites, we compared the structure of isoforms corresponding to those derived from transcripts initiated from internal start sites with PDB proteins using the protein structure predicting algorithm Phyre2 (Mezulis *et al.*, 2015). Corresponding structure predictions were then overlaid on top of each other using separate colors to highlight differences between the two structures. Full-length proteins were also uploaded to NCBI and a domain search was completed using the conserved domain database (CDD). A caveat of this method is that no kinetics can be established, thus the functional efficiency of these predicted proteins remains unknown.

### Motif Analysis

Motif analysis of the Gly-Gly motifs and analysis of the 12mer motifs for pyrimidine and purine frequencies were carried out according to published methods (Grzymski *et al.*, 2014). Gly-Gly motifs were calculated from coding regions broken up by triplets to account for codon frequencies.

## Results and Discussion

### Physiological responses to N-deprivation

Our experimental design was based on previous work examining abrupt N deprivation in *Prochlorococcus* MED4 (Tolonen *et al.*, 2006), facilitating comparisons between previous microarray gene expression results and our analysis of the primary transcriptome. Exponentially growing cells were subjected to acute N stress by resuspension in media with no added N (N-depleted). N-replete media was used as a control. Bulk culture fluorescence values began to diverge three hours after resuspension in N-depleted media; the fluorescence of N-replete cultures continued to increase while the fluorescence of N-depleted cultures decreased (Fig. 1). Maximum PSII photochemical efficiency, Fv/Fm, was used as a measure of the cells’ physiological response to N deprivation (Parkhill *et al.*, 2001). Fv/Fm values for both N-replete and N-deprived cells declined during the first three hours, likely due to the shock of centrifugation and resuspension, but stabilized in the N-replete cells within 12 hours. By contrast, Fv/Fm continued to decline in the N-deprived cells during the course of the experiment (Fig. 1). Following 24 hours in the N-depleted media, the N-deprived cultures had lower bulk fluorescence, fewer cell counts and lower Fv/Fm values compared to N-replete cultures. These results are consistent with previous work (Tolonen *et al.*, 2006) and indicate a significant physiological response of *Prochlorococcus* MED4 to N-deprivation.

**Figure 1.**
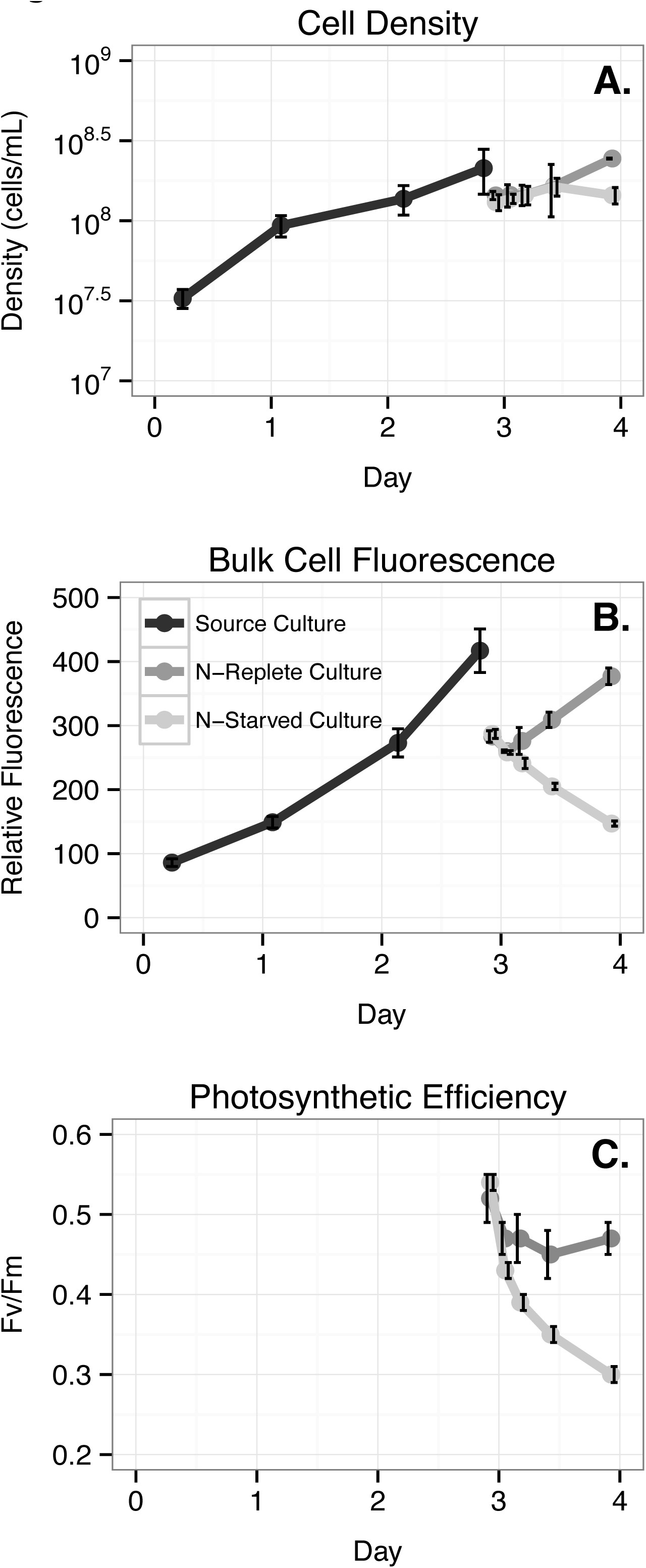
**Physiological changes of *Prochlorococcus* MED4 during N deprivation. A.** Effect of N deprivation on cell concentration. **B.** Effect of N deprivation on bulk culture fluorescence. **C.** Effect of N deprivation on maximum quantum efficiency of photosystem II (Fv/Fm) as measured by fast repetition rate fluorometry. Black lines represent the original triplicate cultures grown in N-replete media; other lines indicate cultures pelleted and resuspended in either N-replete (dark grey line) or N-deficient (light grey line) media. The discontinuity in fluorescence and cell concentration measurements result from incomplete recovery of cells following centrifugation. Error bars represent the standard deviation of 3 biological replicates and are smaller than the symbols when not visible.

### MED4 exhibits a canonical gene expression response to N-deprivation

Using the Ion Torrent reads derived from TEX-treated RNA, we first examined overall gene expression patterns by comparing the transcriptomes of the N-deprived and N-replete cells at 3 different time points following induction of N stress. Overall, the gene expression changes we observed in the primary transcriptome in response to N deprivation were similar to changes previously observed in microarray analysis of total RNA (Tolonen *et al.*, 2006) (Supplementary Text and Supplementary Table S2). We found that the relative changes of differentially expressed transcripts, between N-deprived and N-replete cells, were approximately symmetrical; the number of transcripts demonstrating an increased abundance was similar to the number of transcripts exhibiting a decreased abundance. These data indicate a balanced expression response to N deprivation rather than decreasing transcription overall (Table 1). Down-regulation of a substantial fraction of genes would be a rapid N-saving mechanism and was not the case in *Prochlorococcus* MED4, indicating an active response to N deprivation compared to entry into stasis. Further, the transcriptional response was rapid, with significant changes in transcript abundance apparent within 3 hours of N-deprivation and additional changes noted after 12 and 24 hours (Supplementary Tables S3-S6).

**Table 1.** Differentially Expressed Transcripts between N-deprived vs. N-replete Conditions Per Timepoint (p<0.05)

| Time (hr) | Regulation | Amount of Genes | Differentially |
| --- | --- | --- | --- |
|  |  | Differentially Expressed Genes | Expressed (%) |
| 3 | Up | 62 | 7.21% |
| 12 | Up | 84 | 9.77% |
| 24 | Up | 93 | 10.81% |
| 3 | Down | 28 | 3.26% |
| 12 | Down | 86 | 10.00% |
| 24 | Down | 107 | 12.44% |

Cyanobacteria ‘sense’ N-deprivation by monitoring the C:N balance of the cell and respond to changes in this ratio by activating alternative N assimilation pathways when excess intracellular 2-oxoglutarate accumulates (Ohashi *et al.*, 2011). The global nitrogen stress response regulator, NtcA (Sauer *et al.*, 1999), is responsible for inducing the expression of assimilation pathways for a variety of N compounds. Consistent with previous reports (Tolonen *et al.*, 2006), we found that several members of the NtcA regulon are induced in response to N deprivation in *Prochlorococcus* MED4 (Supplementary Table S5). Members of the NtcA regulon that exhibited increased transcript abundance during N-deprivation included genes encoding the assimilation pathways for urea and cyanate, as well as the glutamine synthase gene *glnA* (Supplementary Tables S3-S6). The NtcA regulon also regulates the urease enzyme and urea transporter enzyme. All subunits of the urease enzyme (*ureA-C*) were co-expressed and showed similar increases in relative abundance at 12 hours post N deprivation. The urea transporter genes (*urtA-E*) were transcribed in a similar manner with the reads gathering at the TSS for *urtA*, the first gene in the operon (Fig. S1). In addition, several high light inducible proteins, known to respond to stress conditions in *Prochlorococcus* (Havaux *et al.*, 2003; Tolonen *et al.*, 2006; Steglich *et al.*, 2008), increased in abundance after 12 hours of N deprivation (Supplementary Table S7). One of these high light inducible proteins, Hli10, is encoded by a gene predicted to be regulated by NtcA (Tolonen *et al.*, 2006).

### Complexity of transcription initiation at variable start sites under N-deprivation

Based on the Ion Torrent data set, gene expression changes in these *Prochlorococcus* MED4 cells exhibited responses similar to previous studies under N-deprivation (Tolonen *et al.*, 2006; Gilbert and Fagan, 2011) suggesting that N savings cannot be achieved by simply downregulating transcription. Thus, to investigate whether *Prochlorococcus* might use other RNA-based regulatory mechanisms to reduce cellular N requirements under N stress, we next examined the primary transcriptome of MED4 following 12 and 24 hours of N depletion. We performed detailed annotation of TSSs after 12 hours post N-depletion due to the strong correlation of transcript patterns at this time point as compared to Tolonen et al. (2006), and confirmed these results with data from 24 hours post N-depletion (Table 2, Supplementary Tables S8-S11).

**Table 2.**
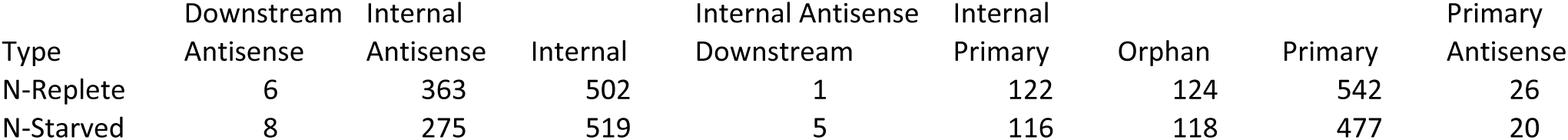
Transcriptional Start Sites By Category

| Type | Downstream<br>Antisense | Internal<br>Antisense | Internal | Internal Antisense<br>Downstream | Internal<br>Primary | Orphan | Primary | Primary<br>Antisense |
| --- | --- | --- | --- | --- | --- | --- | --- | --- |
| N-Replete | 6 | 363 | 502 | 1 | 122 | 124 | 542 | 26 |
| N-Starved | 8 | 275 | 519 | 5 | 116 | 118 | 477 | 20 |

Our detailed annotation of TSSs at 12 and 24 hours following N-depletion identified new internal and antisense TSSs (Table 2, Supplementary Tables S8-S11) throughout the genome that were not previously reported by Voigt et al. (2014) for MED4 grown under N-replete conditions (Supplemental Text, Supplementary Table S12). For the purpose of this study, internal transcription is defined as a verified a saw-tooth pattern of transcript reads mapping downstream of a gene’s annotated start site. The internal transcription ratio is defined as the ratio of read abundance at the internal TSS compared to the abundance of reads at the primary TSS. The internal transcription ratio generally increased under N-deprivation (Fig. 2; Supplementary Table S13), suggesting a specific regulatory response to N stress rather than erroneous transcription given that we did not observe degraded and random read mapping to these genes. Eighty-five genes contained both a primary TSS and internal TSS identified by TSSAR in both the N-depleted and N-replete samples (Supplementary Table S13). Based on a Wilcoxon rank sum test, 64 genes (~74%) demonstrated a significant increase in the internal transcription ratio (p-value < 1x10^−6^ for all 64 genes) in N-depleted samples compared to the N-replete samples, with 13 genes containing multiple internal TSSs (for a total of 81 TSSs in these genes). After 24 hours of N deprivation, transcripts expressed from 72/81 (89%) of these internal TSSs (from 56 genes) were still present, indicating a conserved and continued role in the N stress response.

**Figure 2.**
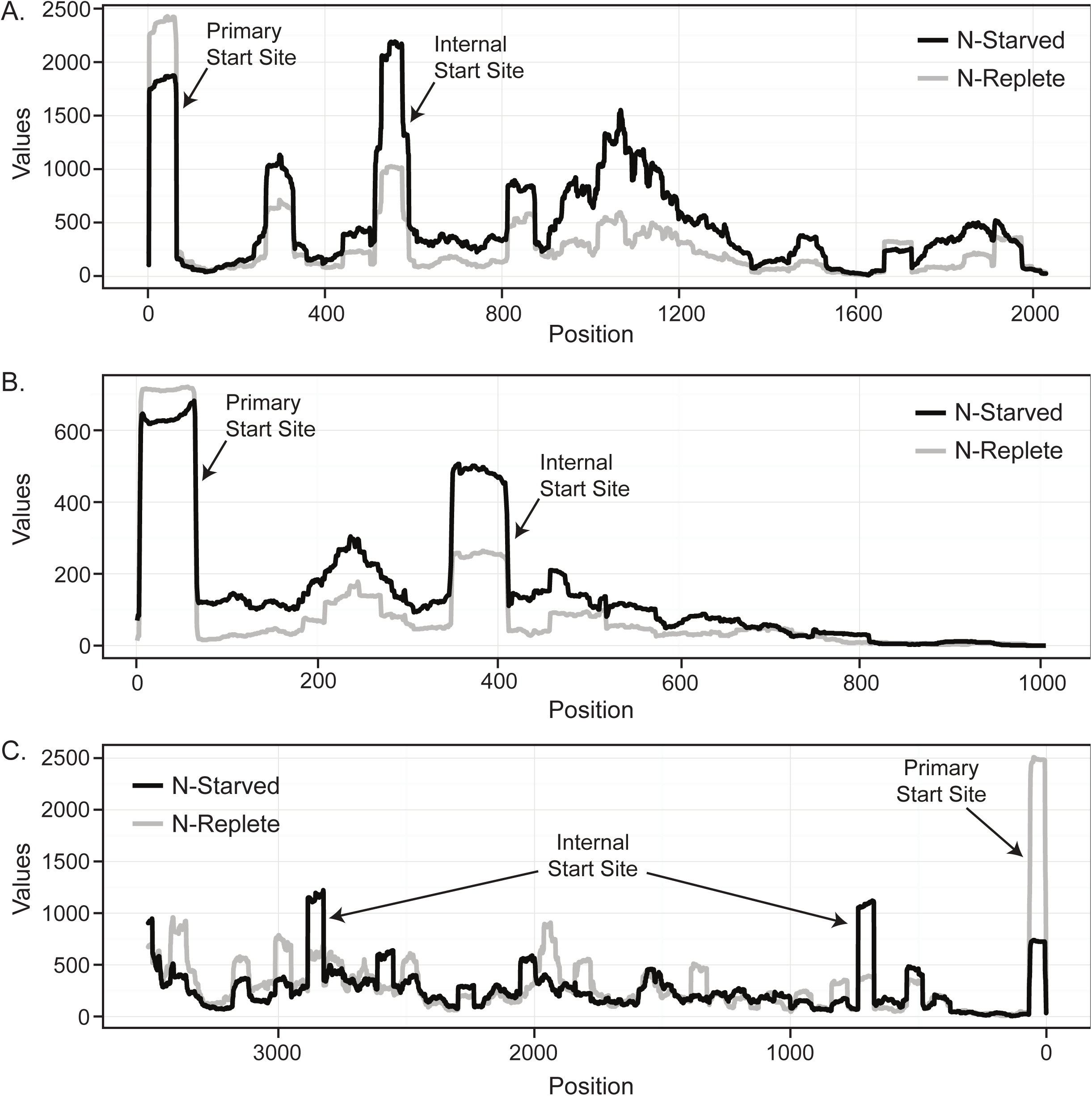
**Primary transcriptome mapping to 3 genes during N deprivation.** Values indicate the number of primary reads mapping to the **A.** PMM0149 (*ndhF*), **B.** PMM0058 (conserved hypothetical protein) and **C.** PMM1485 (*rpoB*) genes. Experimental (red) and control reads (black) were compared at 12 hours post N deprivation. The primary and internal TSSs are marked with arrows in each panel. Each panel represents the full length of the gene, with the x-axis representing the distance from the primary start site.

Transcription from an internal start site, however, was not exclusively a response to nutrient deprivation. We found the internal transcription ratio increased in 13 of the 85 genes (15%) during N-replete conditions compared to N-deprivation, with an additional 8 genes (11%) exhibiting a similar internal transcription ratio in both the N-replete and N-depleted samples (Supplementary Table S13). Irrespective of condition (N-deplete vs. N-replete) 35 genes (~41%) showed a higher abundance of internal TSS reads compared to primary TSS reads, indicating a general increase in internal transcription for those genes. Combined with 187 TSSs from 160 genes that contain only an internal TSS, of which 117 (63%) demonstrated greater internal transcription in N-depleted cells, these results could indicate that some internal TSSs are actually the primary transcription start site which may have been incorrectly called. Consistent with this, Sharma et al. found 18 new gene start sites in their re-annotation of *Helicobacter pylori* using a similar method (Sharma *et al.*, 2010).

On the other hand, we also observed that many of these sites are located far downstream of the annotated start site and well within the expected coding sequence. This suggests that some of these internal TSSs may yield short sense RNAs (intraRNAs) - molecules which have been identified in other studies and are hypothesized to function as truncated alternative mRNAs (Mitschke *et al.*, 2011; Shao *et al.*, 2014). Antisense TSSs were abundant in both N-replete and N-deprived cells, however the relative number of them found throughout the genome was approximately equal in both treatments and cannot be correlated to gene expression changes. These findings further highlight the incredible complexity of transcriptional regulation in *Prochlorococcus.*

### Transcriptional regulation and the N-cost minimization of proteins

Many of the most abundant proteins found within microbes in the Sargasso Sea are transport proteins (Sowell *et al.*, 2008), emphasizing their importance to bacteria living in extremely nutrient depleted environments. NtcA regulation of transporters is known to be an important part of the response of *Prochlorococcus* cells to N-deprivation, as evidenced by a marked increase in transcripts encoding the urea transporter Urt and ammonium transporter Amt1 under these conditions (Tolonen *et al.*, 2006). Based on this, we next focused on the regulation of other genes with possible NtcA binding sites, which could allow them to be regulated as part of the NtcA regulon.

As described above, N-deprivation increased internal transcription in *Prochlorococcus* MED4 (Fig. 2). The intraRNAs initiated from these internal TSSs could have three possible functions: 1) they could be mis-transcribed, degraded or processed RNA with unknown or no function; 2) they could be structural RNAs or encode peptide scaffolds that are used for other regulatory purposes (Lybecker *et al.*, 2014; Shao *et al.*, 2014); or 3) they could be internally transcribed genes that code for functional proteins. While the intraRNAs could arise from transcriptional noise due to spurious promoter-like sequences, and in that case would be non-functional copies of RNA (Lybecker *et al.*, 2014; Shao *et al.*, 2014; Thomason *et al.*, 2015), several genes with internal TSSs have clear regulatory differences in response to N-deprivation (Fig. 2). This has been seen in other cyanobacteria; for example, transcriptional mapping in *Synechocystis* sp. PCC6803 identified abundant short sense transcripts from internal TSSs, which were proposed to yield shorter isoforms of *Synechocystis* 6803 proteins (Mitschke *et al.*, 2011). Random processes would not be expected to show reproducible responses to N-deprivation (i.e., an indication of transcription level regulation). Thus, it is likely these transcripts have a specific physiological role.

We observed that many genes with internal transcription under N-deprived conditions were responsible for important physiological functions. This fact led us to hypothesize that these intraRNAs may encode functional proteins; the shortened RNA transcripts and shortened translated proteins would reduce the overall N requirements of the cell. Given that these cells were already nutrient stressed and exhibiting significantly reduced photosynthesis and growth rates (Fig. 1), the cell might derive a net fitness benefit by expressing a partial protein which required fewer nutrients, as long as that protein retained at least some functionality. To examine this hypothesis, we evaluated the potential effect of the N-terminal truncation on the predicted functional domains of the proteins using NCBI’s conserved domain search (Supplementary Table S14). We also used multiple sequence alignments (MSA) and protein threading to predict the structure of multiple proteins translated from intraRNAs by comparison to known structures (Supplementary Table S14). All of the proteins we examined by protein threading (Supplementary Table S14) contain a possible NtcA binding site upstream of the internal TSS, providing a potential mechanism for internal transcription initiation. We predict that initiating translation at internal start sites would likely have one of three outcomes: 1) proteins that align by MSA to all of the major structural elements of known structures and retain the major carbon backbone; 2) proteins that lack all structural elements or whose structure is clearly incomplete and; 3) proteins where a few specific structural elements or domains are transcribed (Supplementary Table S14).

The cyanate ABC transporter (PMM0370) evidences no structural differences between the full length translated protein and the shortened protein (Fig. 3a, Supplementary Table S14). Although the full diversity of functional cyanate transport proteins with established Protein Data Bank PDB structures have yet to be discovered, our conserved domain analysis results suggest that the shortened protein retains full function of its two predicted domains (Supplementary Table S14). Still, the combined primary transcriptome and modeling results highlight how little we know about many bacterial proteins and further biochemical analysis are warranted to explore how changes in the primary amino acid sequence impact protein function.

**Figure 3.**
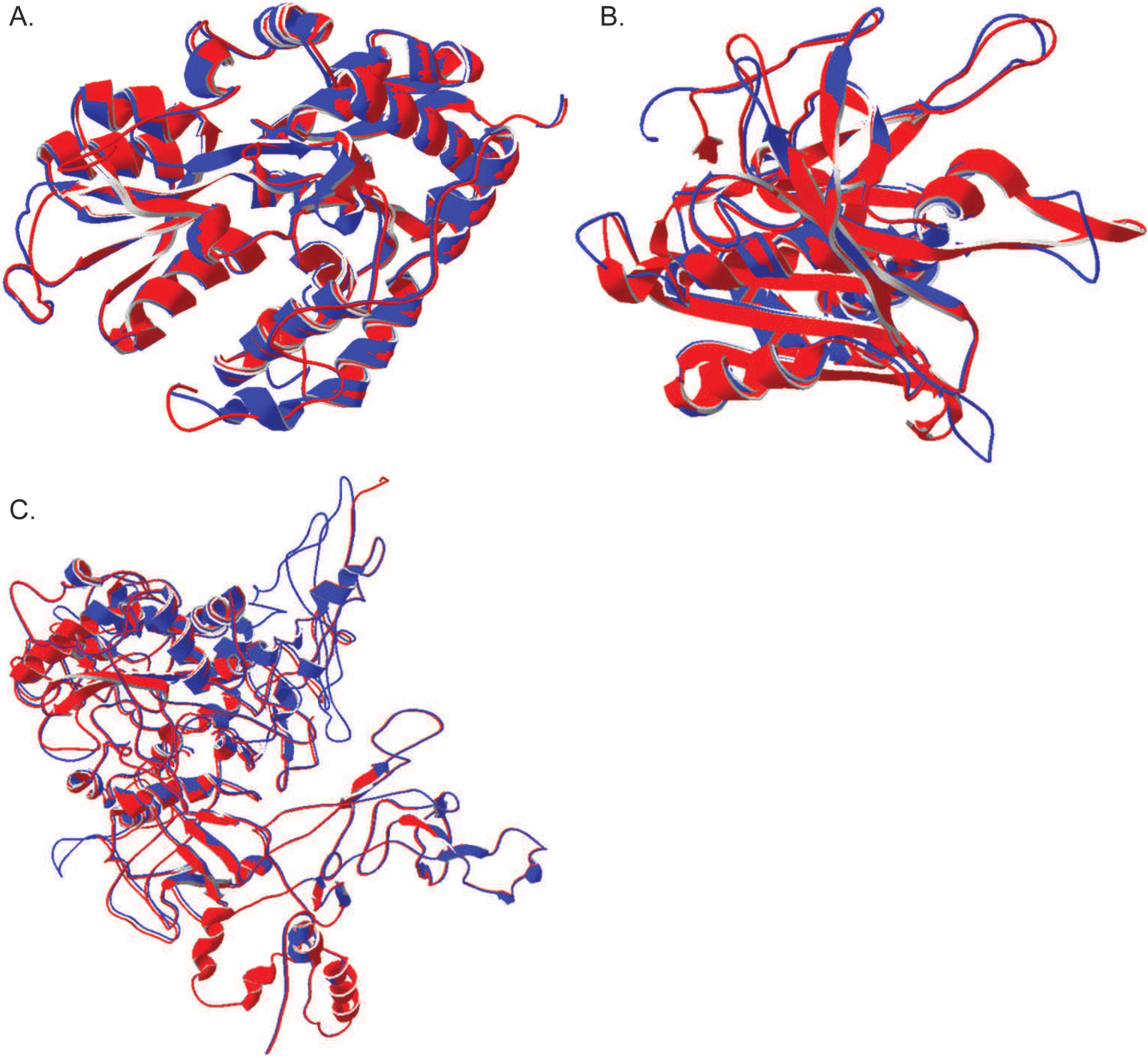
**Structure predictions for internal transcription sites versus PDB structures. A.** Protein structure threading for the cyanate ABC transporter (PDB c2i4cA). There are no major structural differences in the protein threads. Blue and red are the overlapping structure representing the full length transcript (blue) and the corresponding predicted protein from the internal start site in MED4 (red). **B.** As in A, but for the Ferredoxin-NADP reductase protein (PDB C1jb9A). **C.** Protein structure threading for the for RNA polymerase (PDB C3lu0C).

Additionally, a number of other non-transport proteins may have been expressed in shorter forms. For ferredoxin-NADP reductase, there are no discernible changes in the threaded protein structure of the internal transcript, and it likely retains full function of its single predicted domain (Fig. 3b, Supplementary Table S14). RNA polymerase, vital for producing actual transcripts, has an internal TSS whose transcript encodes a protein that aligns to 95% of the sequence from the known PDB structure from *E*. *coli*, although domain function could be slightly modified as the entire protein is predicted as a single domain (Fig. 3c, Supplementary Table S14). It makes sense, physiologically, that under N-stress conditions the protein responsible for producing transcripts is potentially N cost-minimized. If these three proteins were internally translated they would provide 541 mol N / mol translated protein savings relative to the fully translated proteins – approximately 17% savings. The three internal transcripts would provide 4451 mol N/ mol transcript savings compared to the full-length transcripts – a savings of, again, 17%.

In the context of the cell as a whole, transcriptomic N cost minimization is but one of several mechanisms through which *Prochlorococcus* cells adjust their elemental requirements in response to particular stressors such as limitation for key elements such as iron, phosphorus, and nitrogen. For example, in response to iron limitation *Prochlorococcus* expresses the non-iron containing oxidoreductase flavodoxin instead of the iron-sulfur containing ferredoxin (Bibby *et al.*, 2003; Thompson *et al.*, 2011). *Prochlorococcus* also utilizes sulfolipids instead of phospholipids in its cellular membrane in order to decrease phosphorus requirements (Van Mooy *et al.*, 2006). These responses are all based on the utilization of different pathways in order to modify the cell’s elemental requirements. In contrast, transcriptomic cost minimization represents a response to nutrient limitation that depends on structural changes to the mRNA pool and, putatively, the proteome. While genomic N cost minimization, mediated by codon usage and general %GC characteristics, can only change over evolutionary time scales, transcriptomic N cost minimization is a dynamic process which enables the cell to respond to changes in N availability on the order of hours.

### Cost minimized organisms have a higher potential for decreased translation rates

An important consideration regarding both genomic and transcriptomic N cost minimization is that such N savings might only have a small overall impact if protein abundance is not tightly controlled. One strategy for maintaining careful control of protein levels is rapid RNA turnover (Steglich *et al.*, 2010), which allows the cell to quickly adjust mRNA availability in response to stress. *Prochlorococcus,* for example, has a median RNA half-life on the order of 2 minutes, which is twofold faster than that observed in other microorganisms (Steglich *et al.*, 2010). Cells can also improve their control of protein abundance by slowing down translation rates (Sherman and Qian, 2013). While faster growing (r-selected) organisms should experience selective pressures to maximize translation rates in order to support episodes of rapid growth, we hypothesize that slowly growing (k-selected), cost minimized organisms should instead experience selective pressures to minimize translation rates. To explore this hypothesis, we examined the genomes of both oligotrophic k-strategists and copiotrophic r-strategists to look for sequence signatures associated with translation rates.

Shine-Dalgarno motifs (ribosomal binding sites) internal to coding regions have been shown to cause translational pausing and subsequent reductions in growth rate (Li *et al.*, 2012). These signatures are apparent in the observed-to-expected ratio of the frequency of the glycine-glycine motifs in coding regions. Depending on codon usage, Gly-Gly motifs can have high or low affinities to the anti-Shine-Dalgarno sequence found at the 3’ terminus of the 16S rRNA in the ribosome to mediate pausing; such sequences are minimized in fast-growing organisms like *E. coli* (Li *et al.*, 2012). We examined the occurrence of Gly-Gly motifs in the genomes of three bacteria with relatively high maximum growth rates (*Bacillus cereus*, *E. coli* K12 and *Vibrio fisheri*) and four oligotrophic, k-selected marine bacteria with relatively slow maximum growth rates (*Prochlorococcus* MED4, *Prochlorococcus* MIT9313, *Synechococcus* and *Pelagibacter*) (Fig. 4). We found that the genomes of the r-selected organisms exhibited a clear pattern of minimization of Gly-Gly motifs with high affinity to the anti-Shine-Dalgarno sequence, consistent with less pausing and faster protein expression. In oligotrophic k-selected organisms, the pattern is opposite with no apparent deviations in the observed-expected ratios of the motifs; in fact, the motifs with the highest binding affinity to the anti-Shine-Dalgarno sequence were found more often than expected (observed:expected = 1.63) in *Prochlorococcus* MED4. This suggests that selection against motifs that cause translational pausing is weak in oligotrophic k-selected organisms, and therefore that they would be expected to have slower translation rates than the r-strategists. This genomic feature represents yet another property that may contribute to cost minimization in oligotrophic microbes.

**Figure 4.**
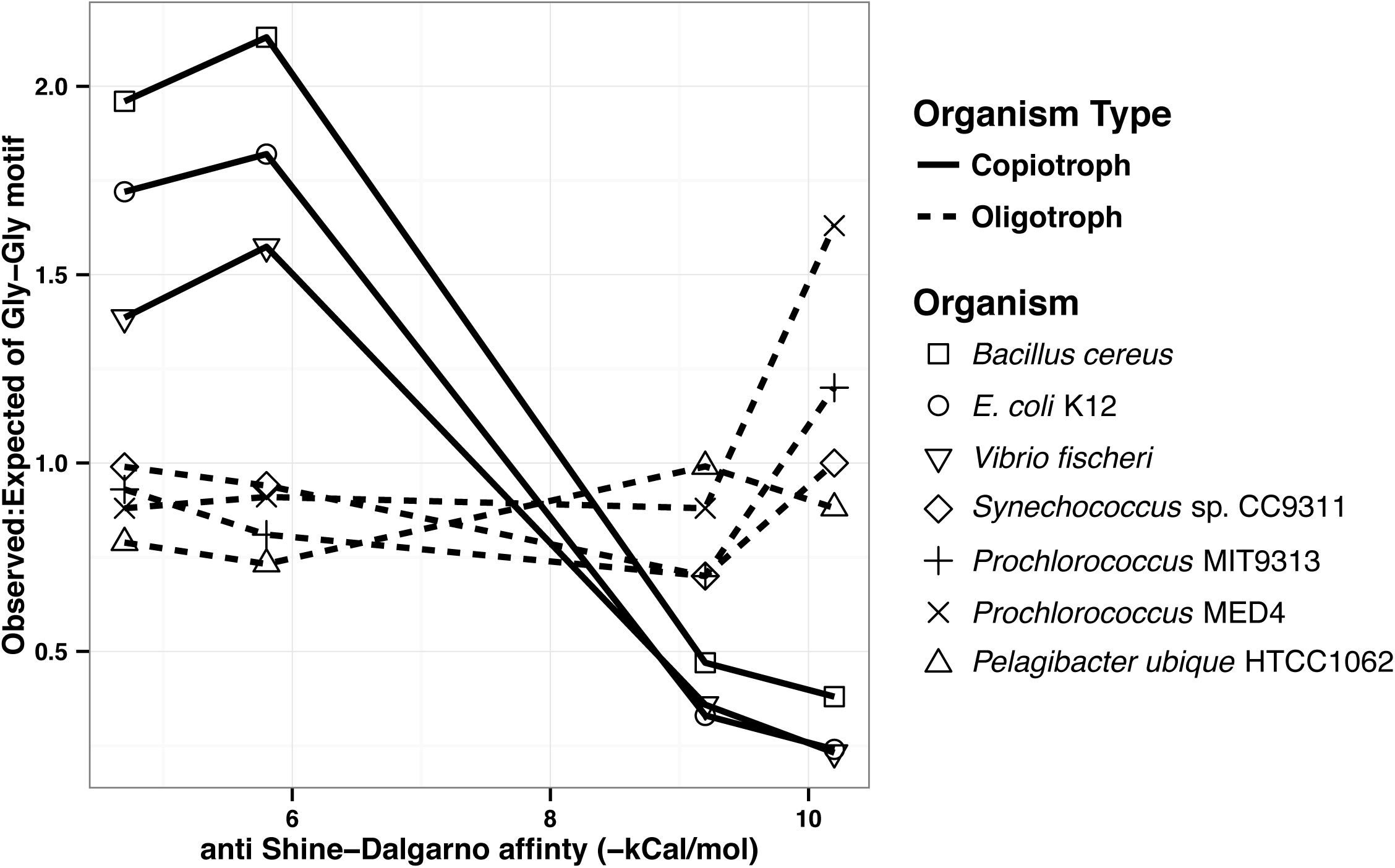
**Frequency of Glycine-Glycine motifs in selected microbial genomes**. Deviations in observed to expected ratios of Gly-Gly motifs, indicative of the potential for ribosomal stalling, are indicated for a set of common copiotrophic r-selected and oligotrophic k-selected organisms.

We propose that there are at least two mechanisms contributing to this weak selection against pausing sequences in slow-growing, oligotrophic bacteria. First, as previously discussed, translational pausing would be beneficial to k-selected organisms by improving the ability of cells to control protein abundance while growing in nutrient limited conditions. Second, it is likely that organisms such as *Prochlorococcus* do not rely on Shine-Dalgarno sequences for translation initiation (Voigt *et al.*, 2014). Instead, based on studies in *Synechococcus,* translation initiation may instead rely on an alternative mechanisms dependent on ribosomal protein S1 (Mutsuda and Sugiura, 2006; Voigt *et al.*, 2014) or the direct binding of a 70S monosome to leaderless mRNA start sites (Moll *et al.*, 2002). This would reduce selection for canonical ribosomal binding sites and, as a less specific mechanism for translation initiation, might provide cells with a way to translate truncated proteins from mRNAs expressed from internal start sites under N stress. These k-selected organisms utilize less regulated and less specific transcription and translation mechanism. As many of these truncated transcripts are not predicted to contain canonical translation initiation sites, S1 binding could offer a mechanistic explanation for the translation of such proteins arising from unexpected transcriptional start sites.

To examine whether ribosomal protein S1-based translation initiation is likely to occur in *Prochlorococcus* MED4, we quantified the relative frequency of 10 and 12mer pyrimidine rich motifs (made up of at least 80% pyrimidines) – sequences which are conductive to S1 binding (Mutsuda and Sugiura, 2006). In connection with the pyrimidine rich regions, we also searched for NtcA binding regions based on the predicted motifs described previously (Su, 2005; Tolonen *et al.*, 2006). Sites were counted as possible NtcA binding sites if they were found less than 100 bp upstream of identified translational start sites, although in some published cases sites can be found much further upstream (Su, 2005). We found abundant 10 and 12mer pyrimidine rich regions directly upstream of annotated translational start sites (Supplementary Table S15). Furthermore, we discovered that many genes had possible NtcA binding sites, which could possibly facilitate NtcA regulation under N stress (Supplementary Table S15). There are also 912 pyrimidine rich motifs found in non-coding regions of the genome (observed:expected = 1.35). Purine rich sequences are concomitantly under-observed and, given that there are no G+C biases in purine and pyrimidine motifs, these observations are consistent with the hypothesis that S1 translation initiation can occur both in 5’ untranslated regions and within sequences downstream of traditional start sites.

These data suggest that protein S1 could associate with these pyrimidine rich sequences and mediate translation of proteins expressed from the annotated primary start site or, as suggested by our transcriptome data under N deprivation conditions, from internally transcribed start sites. Leaderless mRNAs are likely present within *Prochlorococcus* MED4, requiring an alternate method of translation initiation. In these cases, mRNAs lacking a 5’ UTR directly bind 70S monosomes, thus initiating translation (Moll *et al.*, 2002; Voigt *et al.*, 2014). Our data, consistent with previous studies on *Prochlorococcus* MED4 (Voigt *et al.*, 2014), suggest that 6-8% of all primary TSSs are found within 10nt of translation initiation sites and thus potentially initiated by 70S monosomes.

## Conclusions

In summary, we have shown that a variety of structural changes occur within the *Prochlorococcus* transcriptome in response to N deprivation, and that these changes likely contribute to the ability of this organism to minimize the amount of N required in its proteome. Specifically, *Prochlorococcus* increased internal transcription for several genes during N stress conditions, which should in some cases produce shortened versions of enzymes that likely retain at least partial functionality and require fewer N atoms. Proteomic confirmation of the translation of the shortened peptides, as well as biochemical characterization, will be necessary to understand their abundance and function relative to their full-length versions. We also propose that *Prochlorococcus* may have relatively slow translation rates which, in conjunction with short RNA half-lives, allows them to control protein abundances, reducing cellular nitrogen requirements. Although the concept of cost minimization has heretofore been considered to function on evolutionary time scales (through selection on genomic codon usage to reduce nutrient requirements), these results show that cost minimization can encompass physiological mechanisms as well – through dynamic structural changes to the transcriptome that should result in a proteome requiring fewer N atoms.

## Acknowledgements

We thank Alexis Yelton (MIT) and Julie Miller (MIT) for assistance with sampling. We also thank Dr. David Vuono (DRI) for his contributions in the preparation of the manuscript. This work was funded in part by the Gordon and Betty Moore Foundation through Grant GBMF495 to SWC, by a grant from the Simons Foundation (SCOPE award ID 329108 to SWC), and by the National Science Foundation (MCB-1244630 to JJG and OCE-1153588 and DBI-0424599 to SWC). ACR was supported by a HHMI International Student Research Fellowship. This article is a contribution from the Simons Collaboration on Ocean Processes and Ecology (SCOPE).

